# Efficient and accurate prime editing strategy to correct genetic alterations in hiPSC using single EF-1alpha driven all-in-one plasmids

**DOI:** 10.1101/2022.05.04.490422

**Authors:** Wout J. Weuring, N Dirkx, E De Vriendt, N Smal, J van de Vondervoort, Ruben van ’t Slot, M Koetsier, S Weckhuysen, Bobby PC Koeleman

## Abstract

Prime editing (PE) is currently the most effective and versatile technology to make targeted alterations in the genome. Several improvements to the PE machinery have recently been made, and have been tested in a range of model systems, including immortalized cell lines, stem-cells and animal models. While nick RNA (ncRNA)-dependent PE systems like PE3 and PE5 are currently considered to be the most effective, they come with undesired indels or SNVs at the edit locus. Here, we aimed to improve ncRNA-independent systems PE2 and PE4max by generating novel all-in-one (pAIO) plasmids, driven by a tissue-broad EF-1alpha promoter, that is especially suitable for human iPSC models, and linked to a GFP tag for fluorescent based sorting. These novel pAIO systems effectively corrected mutations observed in patients suffering from epileptic encephalopathy, including a truncating *SCN1A* R612* variant in HEK293T cells and a gain-of-function *KCNQ2* R201C variant in patient-derived hiPSC, with edit efficiency up to 50%. We also show that introducing additional silent PAM-removing mutations can negatively influence edit efficiency. Finally, we observed an absence of genome-wide PE off-target effects at pegRNA homology sites. Taken together, our study shows an improved efficacy and accuracy of EF-1alpha driven ncRNA-independent pAIO PE plasmids in hiPSC.

## Introduction

CRISPR/Cas9 was first used for genome editing in eukaryotic cells in 2013 ^[1]^, after which it rapidly evolved into a genome editing tool that has revolutionized the field of biology as a whole. Not only did it lead to an unprecedented increase in our capacity to study gene function and the effects of genetic variation on health and disease. It also brought us closer to the implementation of true precision medicine: the ability to apply treatment strategies specifically targeting the cause of disease; the underlying gene mutation. Prime editing (PE), the most recent gene editing tool in the field, uses a Cas9 nickase (Cas9n) protein fused to a reverse transcriptase (RT), and a PE guide RNA (pegRNA) consisting of a guide, a primer binding site, and RT template containing the desired edit ^[2]^. PE is the first gene edit system that can introduce all four possible base pair (bp) transversions, small insertions and deletions. It also has lower off-target effects and a higher edit-to-indel ratio compared to previous Cas9n editors. The first-generation of PE consist of PE2 (pegRNA + Cas9n-RT) and PE3 (pegRNA + Cas9n-RT + additional nicking RNA (ncRNA)). PE3 generates a second single-stranded break via ncRNA activity and enhances editing efficiencies by 1.5 to 4.2 fold compared to PE2 in immortalized cell lines, it however also increases the number of undesired indels and single nucleotide variants (SNV) ^[2]^. The second generation PE systems have a human codon-optimized RT, a further optimized Cas9n and chemically modified pegRNAs that increase stability. PE2 and PE3 systems that have these enhancements can furthermore be co-expressed with MutL homolog 1 (MLH1dn), which is a DNA mismatch repair (MMR)-inhibiting protein, and are then named PE4max and PE5max, respectively ^[3]^.

When PE is applied to more advanced human *in-vitro* model systems such as human induced pluripotent stem cells (hiPSC) or liver organoid cells, edit efficiencies decrease significantly and are between 0 and 0.83% if no additional selection procedures are applied ^[4][5]^. hiPSC-derived tissues are one of the most frequently used models to study disease mechanism or treatment strategies for genetic disorders nowadays. Where it was initially a common practice to use patient- and healthy control-derived iPSC lines as an experimental setup, it is nowadays preferred to either introduce disease relevant mutations in a control background or to correct mutations in patient-derived iPSCs to generate isogenic lines. This also allows the study of rare pathogenic variants or variants of unknown significance, when patient material is not available. However, introducing or correcting mutations is still highly laborious and costly, creating a bottleneck for more high-throughput projects. Recently, an all-in-one plasmid (pAIO) was published that contains the complete PE3 system (PEA1) thus facilitating transfection of stem cells with all the necessary constructs at once ^[6]^. The pAIO-PE3 plasmid, combined with a puromycin selection showed a significant increase compared to the regular PE3 system in mouse stem embryonic stem cells, with edit efficiencies ranging between 1.7% and 85%. In line with other ncRNA-dependent PE systems, pAIO-PE3 activity however also resulted in a high rate of indel/SNVgeneration for most loci, occurring in up to 90% of the clones ^[6]^. Since PE3 systems are accompanied by higher undesired indels/SNVs, improved versions of ncRNA-independent PE systems would be safer alternatives for gene-editing in iPSC-derived models. Furthermore, for optimal use in human iPSC disease models, elongation factor 1alpha (EF-1alpha) represents a more tissue-broad promoter that can drive stable expression during iPSC differentiation ^[7][8]^.

For these reasons, we generated an pAIO-PE2-GFP and pAIO-PE4max plasmid, driven by the EF-1alpha promoter as an improvement to current cytomegalovirus (CMV)-driven prime editors and improved edit efficiencies by adding a cell sorting step.

## MATERIAL AND METHODS

### PegRNA cloning for PE3, PE2max and PE4max

pegRNAs and nicking RNAs were cloned into *AAV-U6-sgRNA-hSyn-mCherry* (Addgene #87916). The pegRNAs and nicking RNAs were designed using PegFinder ^[9]^. 100uM single-stranded DNA oligos were ordered via IDT in MilliQ. Oligos were annealed by mixing 10 uL of each oligo in 20 µL of 2X annealing buffer (20 mM Tris pH 7.5, 100 mM NaCl and 20 mM EDTA) and ran in the following PCR program; 5 minutes on 95°C, start on 95°C and -2°C/second for 4 seconds, start on 85°C and - 0.1°C/second for 599 seconds and incubation on 4°C. The AAV-U6-mCherry plasmid was linearized by SapI following manufacturer’ s instructions. Ligation of the oligos was performed by T4 DNA Ligase (NEB M0202L) following manufacturer’ s protocol and transformed using *TOP10* competent cells. Plasmids were purified using Qiagen mini or midiprep kits and verified by Sanger Sequencing. For oligo sequences see **Supplementary data S1**

### Generation of pLV-EF1a Prime Edit plasmids

pLV-EF1a-PE2-P2A-GFP was generated by cloning the original pCMV-PE2-P2A-GFP insert (Addgene #132776) into an EF-1alpha driven lentiviral backbone by Signagen Inc (from here on called pLV-PE2). DNA oligos were designed for introduction of new restriction sites in pLV-PE2. Single-stranded DNA oligos were annealed following previous described protocol. Digestion of the backbone was performed by ClaI (NEB #R0197L) according to manufacturer’ s instructions. Ligation of the annealed oligos in the linear backbone was performed by T4 DNA Ligase. pLV-EF1a-PEmax-P2A-hMLH1-dn was generated from the pLV-PE backbone. The pCMV-PE4max backbone was linearized by AgeI (NEB #R3552L) and MluI restriction sites were introduced via molecular gene cloning according to above methods with the exception that *Stbl3* and not *TOP10* competent cells were used. The pCMV-PE4max with MluI restriction site and the pLV-PE plasmid were both digested by MluI (NEB #R0198L) and NotI (NEB #R0189L). PEmax-P2A-hMLH1-dn insert was ligated in the pLV-EF1alpha backbone and transformed in *Stbl3* competent cells. pLV-EF1a-PE2-P2A-GFP (#184445) and pLV-EF1a-PEmax-P2A-hMLH1-dn (#184444) are available on Addgene after publication. Plasmids were verified using Sanger Sequencing. For oligo sequences see **Supplementary data S1**

### pegRNA cloning and generation of pAIO-PE2-GFP and pAIO-PE4max

PegRNAs were ordered as 500ng gBlock (double stranded DNA fragments) via IDT and resuspended in 10µL MilliQ prior to cloning in the dual PaqCI (NEB #R0745L) site of pLV-EF1a-PE2-P2A-GFP (from here on called pAIO-PE2-GFP) or pLV-EF1a-PEmax-P2A-hMLH1-dn (from here on called pAIO-PE4max). Digested plasmids and gBlock DNA fragments were purified by Wizard SV Gel and PCR clean-up system (Promega) and transformed in *Stbl3* competent cells. pAIO-PE2-GFP:KCNQ2-C201R (#185060) and pAIO-PE2-GFP:KCNQ2-C201R_pp (#185061) are available on Addgene after publication. Plasmids were verified using Sanger Sequencing. For oligo sequences see **Supplementary data S1**

### Prime editing transfections in HEK293T cells

24 hours prior to transfection, HEK293T cells were seeded to yield approximately 50% confluency at day 1 in 6 wells plates. For PE3, three different plasmids were transfected simultaneously in HEK293T cells; 1 µg of pCMV-PE2-P2A-GFP, 0.5 µg of pegRNA plasmid and 0.5 µg of nicking RNA were combined with 50 uL OptiMEM and 4 uL Lipofectamine2000 and added to the cells. For PE2max (Addgene #174820) and PE4max (Addgene #174828), 1.4 µg editor and 0.6 µg of pegRNA was used. Genomic DNA extraction was performed 72 hours after transfection using DNeasy Blood & Tissue Kit from Qiagen. In the case of a second transfection experiment, approximately 20% of the cells remained in culture and transfected when confluency reached 50% again. The remaining cells were used for gDNA extraction. The edit locus was PCR amplified using TaqGold Polymerase and the PCR product was purified by Wizard SV Gel and PCR clean-up kit followed by Sanger Sequencing.

### Whole exome sequencing on edited HEK293T

Whole exome sequencing was performed on DNA from HEK293T. Methods and the script used are listed here: https://github.com/UMCUGenetics/DxNextflowWES and the IGV alignment is added as **Supplementary data S3**.

### Lentivirus production

Lentivirus (LV) was produced by transfection of transfer and packaging plasmids in HEK293T cells. The following plasmids were used in a 4:2:1:1 (ug) ratio; transfer plasmid (pAIO): pMDLg/pRRe (or psPAX2-D64 for integrase deficient lentivirus); pRSV-rev; pMD2.g. pDNA was mixed in 200 uL OptiMEM and combined with 24 uL PEI (1 mg/mL). The DNA:PEI complexes were added to the cells after 20 minutes incubation at room temperature and medium was replaced at approximately 18 hours. LV or integrase-deficient LV (IDLV) particles were harvested using the Lenti-X-concentrator (comp/code) 72 hours after transfection according to manufacturer’ s protocol. The titer of harvested lentivirus was established using the Lenti-X RT-qPCR Titration Kit (Takara Bio, Cat. no. 631235). A representative calibration plot and LV tittering of pAIO is added as **Supplementary data S4**.

### hiPSC maintenance

The hiPSC line harboring the R201C *KCNQ2* variant was obtained by reprogramming patient PBMCs. hiPSCs were maintained in StemFlex medium (Thermo Fisher Scientific, A3349401) on Geltrex (Thermo fisher Scientific, A1413302) coated plates at 37°C and 5% CO2. Cells were passaged with ReleSR (Stemcell Technologies, 05872) when they reached 70-80% confluency. The hiPSC line was tested for pluripotency markers (Nanog, Sox2 and Oct4) using qPCR, and showed no abnormalities on CNV analysis.

### Prime editing nucleofection of hiPSC

hiPSC were dissociated using Accutase (Sigma, A6964). Cell pellets of 1E6 cells were resuspended with 20ul supplemented nucleofector solution of the P3 Primary Cell 4D-nucleofector® X Kit S (Lonza, V4XP-3032) and 2ug plasmid (1ug/ul), and nucleofected with the 4D-Nucleofector Core and X Unit (Lonza), with program CB-150. 10 min after nucleofection, cells were plated in StemFlex media supplemented with 10uM Y-27632 rho kinase inhibitor (Tocris, 1254). Genomic DNA extraction was performed 72 hours after transfection, unless stated otherwise, using Macherey-Nagel NucleoSpin DNA mini kit (Macherey-Nagel, 740952) according to manufacturer’ s protocol. The edit locus was PCR amplified using Titanium® Taq DNA Polymerase (Takara, 639208) followed by Sanger Sequencing.

### Flow cytometry analysis and fluorescence-activated cell sorting (FACS) of iPSC nucleofected with pAIO-PE2-GFP

48 hours after nucleofection, hiPSCs were dissociated using Accutase. The pellet was resuspended in 1ml FACS buffer (PBS with 2% Fetal bovin serum), with 1000X Viability Dye eFluor® 780 (Thermo Fisher Sci., 65-0865), and incubated for 20 min on ice. After incubation, 10ml FACS buffer was added, and cells were centrifuged for 3min at 300g to wash the cells. The cell pellet was resuspended in 2 ml FACS buffer and ran through a 30um filter. GFP+ signal was measured using the MACSQuant® Analyzer. Cells for sorting were added to the MACSQuant® sorting cartridge (Miltenyi Biotec). Gating hierarchies were constructed using MACSQuant Tyto software before sorting. Cell debris, doublets, and dead cells were gated out. GFP+ signal was divided in two (high and low GFP signal). After sorting cells of the lower GFP signal, the positively sorted cells were collected from the cartridge and seeded in StemFlex media supplemented with 10uM Y-27632 rho kinase inhibitor, the negatively sorted cells were reloaded, and the sorting gate was changed to a higher GFP+ signal.

### Subcloning prime edited iPSC line

Subcloning of the edited iPSC lines to generate isogenic clones, was performed by seeding 500 single iPSCs in one well of a 6 well plate coated with Geltrex, in StemFlex media supplemented with CloneR™ (stem cell technologies, 05888). Three hours after seeding, cells were placed in the Incuctye® Live Cell Analysis System and imaged every 3 hours for 7 to 10 days to track if colonies originated from 1 single cell. Cells were kept in CloneR supplement media for 4 days, whereafter the CloneR supplement was removed. Between day 7 and day 10, colonies originating from one single cell were picked and placed in an individual well of a 24 well plated coated with Geltrex and StemFlex media. Clonal lines were split into two 12 wells; one for expansion and one for DNA isolation to check the absence or presence of the R201C mutation.

### Determination of edit efficiency and off-target activity

Sanger files (.ab1) were uploaded in EditR for determination of edit efficiency and unintended SNVs or indels. Sanger files for **Figure 1D** is added as **Supplementary data S2**.

**Figure 1.**
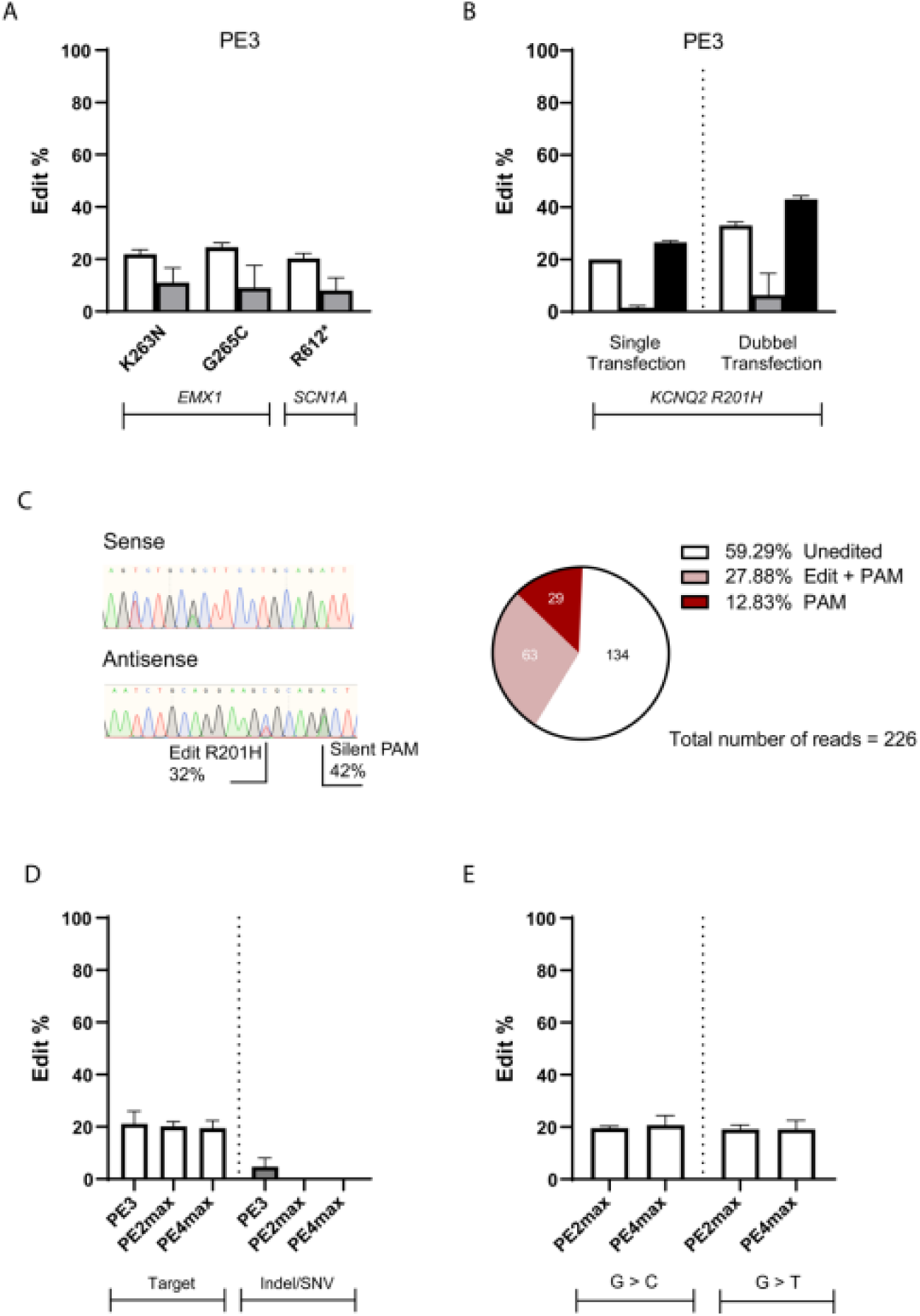
Introduction of DEE mutations using PE3, PE2max and PE4max. White bars represent editing of the target of interest, grey bars represent presence of undesired SNVs or indels at the edit locus, and black bars represent editing of the silent-PAM removing mutation. A) Validation of PE3 editing in EMX1 (K230N. G232C) and SCN1A (R612*). B) PE3 introduction of KCNQ2 R201H mutation with an addition silent-PAM removing mutation. C) Sanger and WES comparison of R201H double transfected PE3 sample. D) Comparison of ncRNA-dependent and independent editing systems using 3 different pegRNA. E) Efficiency ofGtoC conversion compared to G to T conversion in SCN1A at position c.1863-1865. The mean ± SD of all individual values of n = 3 to 6 independent replicates are shown.

### Whole genome sequencing analysis on edited hiPSC

DNA was extracted from two sorted clones and the naïve parental hiPSC line. Samples were submitted to BGI Denmark for WGS with standard sequencing coverage of 30X/90Gb data. Reads were aligned to human genome build hg38 using BWA. Single-nucleotide variants and small insertion-deletions were called using both GATK and Strelka variant callers. Novel variants in the predicted off-target regions of the corrected hiPSC clones were identified as any variant within 250 bp of a predicted off-target site with a mismatch threshold of 4 that was called as reference in the parent clone by both GATK and strelka and as a variant in one of the two corrected clones by both callers (with coverage > 8 and genoqual > 25). Novel variants in the whole genome were further filtered on having no supporting reads in the parent clone, a GATK mapping quality > 50, mapping quality rank score > -2.5 and genotype quality > 70 in both the parent and corrected clone. Variants were annotated using GenomeComb.

### qPCR pluripotency markers iPSC

Total RNA of five corrected hiPSC clones and the naïve parental iPSC line was isolated using the Macherey-Nagel NucleoSpin RNA mini kit (Macherey-Nagel, 740955) according to manufacturer’ s protocol. cDNA was reverse-transcribed from total RNA using the iScript cDNA Synthesis Kit (Bio-Rad Laboratories, 1708890). qPCR reactions were done in triplicate with SYBR Green Real-Time PCR master mixes (Applied Biosystems, 4309155). Expression levels were normalized to GAPDH and the ΔΔCt method was used to determine the relative levels of mRNA expression (**Supplementary data S6)**.

### Multiplex Amplicon Quantification

In order to determine whether the recurrent chr20q11.21 duplication ^[10][11]^ appeared in the corrected clonal lines, we screened our corrected clones with the in-house developed Multiplex Amplicon Quantification (MAQ) technique (Agilent) consisting of a multiplex PCR amplification of fluorescently labeled target and control amplicons, followed by fragment analysis. The assay contains six target amplicons located in and around the chr20q11.21 region, and five control amplicons located at randomly selected genomic positions outside the chr20q11.21 region and other known CNVs. These 11 amplicons were PCR-amplified in a single reaction containing 20 ng of genomic DNA. Peak areas of the target amplicons were normalized to those of the control amplicons. Comparison of normalized peak areas between clonal lines and references resulted in a dosage quotient (DQ) for each target amplicon, calculated by the MAQ software (MAQ-S) package (Agilent). DQ values above 1,25 were considered indicative for a duplication (**Supplementary data S5)**.

## RESULTS

### Introduction of developmental epileptic encephalopathy variants in HEK293T using multi-plasmid PE systems

Pathogenic variants in *SCN1A* and *KCNQ2* are both associated with Developmental Epileptic Encephalopathies (DEE), a group of severe neurological syndromes with an early age of onset and severe prognosis. *KCNQ2* R201H is a Gain of Function variant that leads to a severe neonatal-onset encephalopathy without neonatal seizures but prominent startle-like myoclonus, and a burst-suppression EEG pattern ^[12]^. The SCN1A R612* truncating variant was previously described in an individual with a DEE prototype Dravet Syndrome and gave rise to a severe seizure phenotype that resulted in acute encephalopathy leading to death ^[13]^.

As a proof of principle, we replicated an experiment from the original PE publication in HEK293T cells using PE3 to introduce mutations in *EMX1* (K263N and G265C) ^[2]^. We then showed we could reach comparable edit levels (∼20%) when introducing a pathogenic variant in *SCN1A* (R612*) **[Figure 1A**]. As expected for the ncRNA-depedent PE3 system, all conditions showed unintentend Indels/SNVs.

Next, the *KCNQ2* R201H mutation was introduced with an additional silent protospacer adjacent motif (PAM)-removing mutation. After the first round of transfection with PE3, both the edit and the silent PAM-removing mutation were detected. Interestingly, the edit levels of silent PAM-removing mutation exceeded that of R201H **[Figure 1B]**. Another round of PE3 increased the edit levels for both R201H and silent PAM. We performed whole exome sequencing to validate edit efficiencies and to determine whether the exceeded rate of PAM removal was the result of a biased PCR amplification and Sanger sequencing. WES analysis confirmed that approximately 28% of the reads carry both R201H and PAM removal, but another 12% carried only the silent PAM mutation yielding approximately 40% of the reads with a PAM mutation [**Figure 1B & Supplementary data S3 / BAM file**]. These findings highlight that Sanger sequencing gives comparable results as NGS for PE3 experiments for R201H and PAM removal in HEK293T cells. Furthermore, introduction of an addition silent PAM-removing mutation can be favored over the intended edit. Cells with only the PAM-removal edit can not be edited anymore as the PE system is unable to nick the target DNA, therefore, reducing edit efficiencies.

Next, we compared the edit efficiency of the second generation, PE2max and PE4max multi-plasmid systems to PE3 for 3 pegRNAs in 2 different loci. All three editing systems are approximately equally effective, however, unintentend Indels/SNVs were only detected for PE3, highlighting the challenge of ncRNA-dependent PE [**Figure 1C**]. Due to the PE4max ability to inhibite MMR, it was previously shown to have higher edit efficiencies compared to PE2max. Interestingly, we did not observe this for our edited loci. Because C to G conversions can be more efficiently introduced as they evade MMR ^[3]^, we also validated if changing a NGG motif in *SCN1A* at position c.1863-1865 (NM_001165963.4) to NCG (p.L623V) would be more efficient compared to NTG (p.L623M). No increased efficiency was observed for L623V compared to L623M using PE2max. Both observations can be explained by the MMR deficiency of HEK293T cell, which limits the MMR-inhibition effect of PE4max and the advantage of specific base changes **[Figure 1D**].

### Generation of all-in-one EF-1alpha driven PE systems

As an improvement to current CMV-driven PE systems that require several plasmids [**Figure 2A**], we generated two novel all-in-one EF-1alpha-driven PE systems on a lentiviral backbone using molecular gene cloning. The tissue-broad EF-1alpha promoter is known for its high activity and stability in iPSCs over time, and is also described as one of the most stable promoters during iPSC differentiation ^[7][8]^. Because ncRNA-based PE systems typically lead to additional unintended SNVs and indels, we used the PE2-P2A-GFP editor ^[2]^ with a pegRNA to yield pAIO-PE2-GFP, and one of the most recent editors PEmax-MHL1dn ^[3]^ with a pegRNA to generate pAIO-PE4max [**Figure 2B]**.

**Figure 2.**
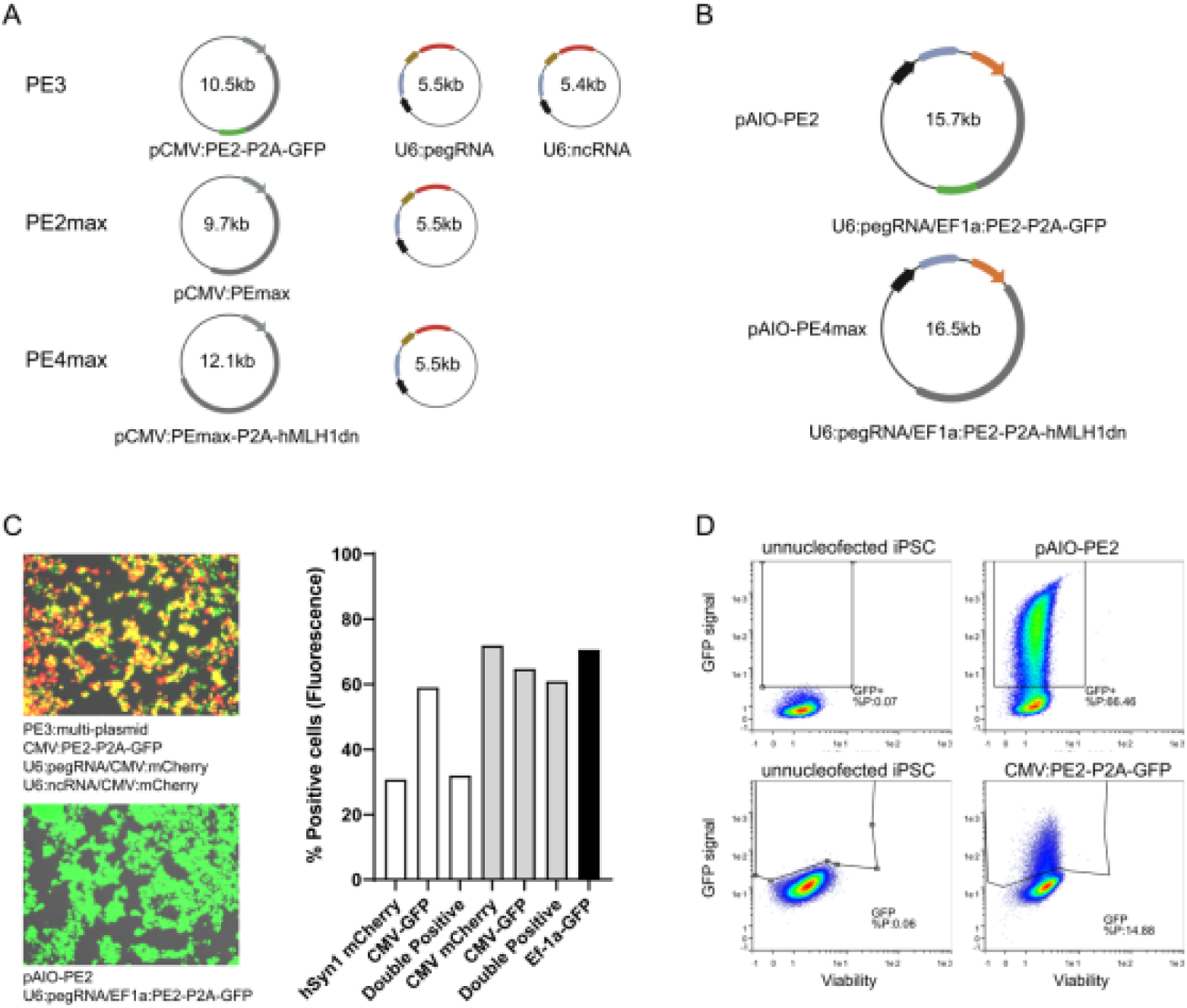
Generation of all-in-one ncRNA-independent PE systems A) Overview of CMV-driven multiplasmid PE systems B) Overview of the two pAIO plasmids generated for this study. Plasmids are abbreviated to pAIO-PE2 and pAIO-PE4max. Grey arrow = CMV, Black arrow = U6. Brown arrow = hSyn1 Orange arrow = EF-1alpha. C) Comparison of multi-plasmid PE3 using two different promotors, versus pAIO-PE2-GFP delivery in HEK293T cells. The fraction of total mCherry, GFP and double positive (mCherry+GFP) cells are plotted. The black bar represents pAIO-PE2-GFP delivery. D) Delivery of PE3 and pAIO-PE2-GFP plasmids to iPSCs measured via flow cytometry and compared to unnucleofected iPSCs.

Because the size of a plasmid influences transfection efficiency, we validated the delivery efficiency of our AIO plasmids which are 1.5 to 3 times larger than the independent PE plasmids [**Figure 2B**]. We transfected HEK293T cells with our novel pAIO-PE2-GFP and compared it to multiplasmid PE3 transfections using the original pCMV:PE2-P2A-GFP and mCherry-tagged pegRNA and ncRNA plasmids. As expected for the PE3 system, a subset of cells underwent partial delivery shown by only expressing mCherry or GFP [**Figure 2C**]. Delivery with pAIO-PE2-GFP showed similar expression to that of multi-plasmid PE3 in HEK293T, proving efficient delivery of our bigger plasmids (**Figure 2C**). Furthermore, use of a suboptimal promoter in the multi-plasmid system, shown here as the hSyn1 promoter in HEK293T cells, significantly decreased the amount of double positive transfected cells compared to the CMV-driven multi-plasmid experiment (**Figure 2C** right).

Next, we validated the delivery of the pAIO-PE2-GFP to PE3 in iPSCs via flow cytometry. In contrast to our results in HEK293T cells, FACS analysis of the GFP positive population showed a 4-fold higher expression for the pAIO-PE2-GFP system compared to the CMV driven multi-plasmidPE3 system (66% vs 15%, respectively) [**Figure 2D**]. Together, these data show the effective delivery of our AIO-PE system in HEK293T cells and iPSC, and highlight the importance of an optimal promoter and a single source of the PE machinery for transfection of iPSC.

### Removal of homozygous *SCN1A* R612* variant using pAIO systems in HEK293T

Using PE3, we generated a homozygous R612* HEK293T cell line via single cell sorting and clonal expansion [data not shown]. To validate if pAIO plasmids for the correction of this mutation were functional, HEK293T^R612*/R612*^ cells were transfected with pAIO-PE2-GFP and pAIO-PE4max and compared to PE3. After one transfection, pAIO-PE2-GFP, pAIO-PE4max and PE3 repaired the *SCN1A* R612* mutation on average for 31%, 33% and 29.5% of the cell population, respectively, validating that pAIO plasmids are functional. No undesired SNVs or indels were detected in pAIO-transfected cells, in contrast to those transfected with PE3 [**Figure 3**].

**Figure 3.**
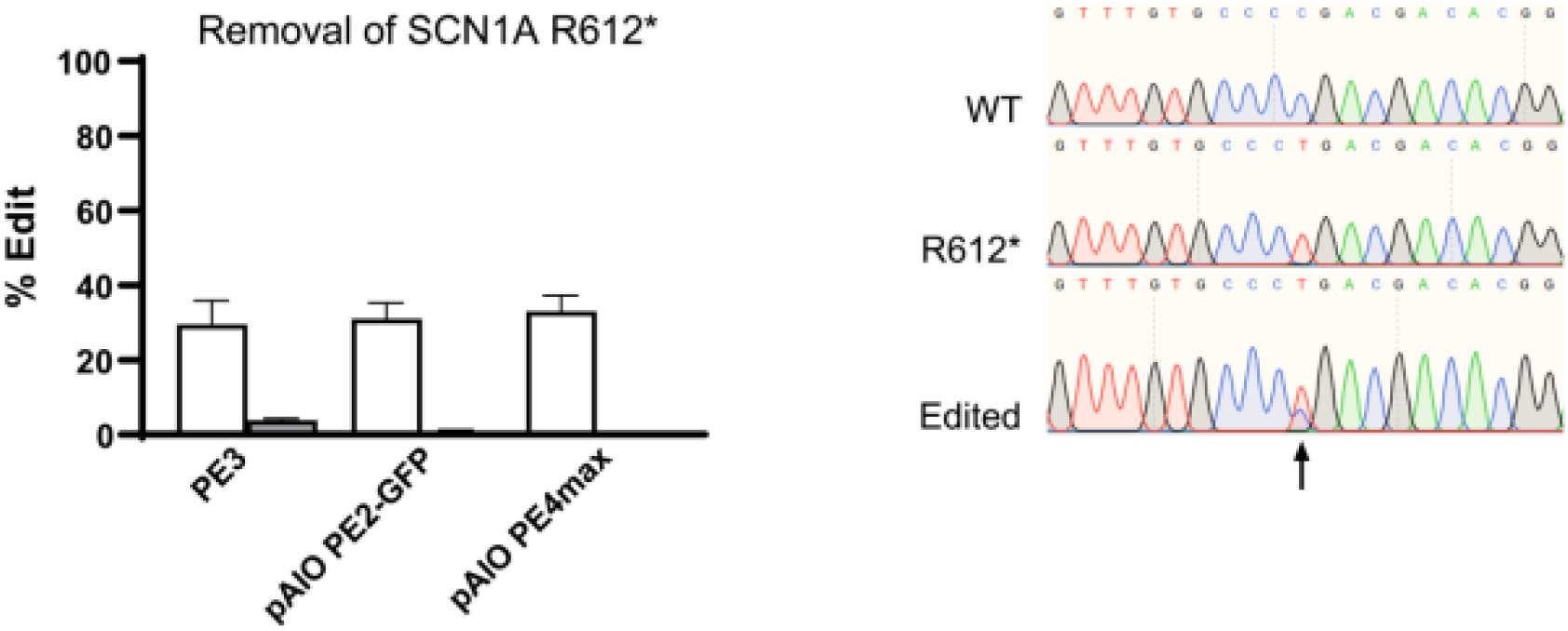
Correction of SCN1A R612* in HEK293T cells. Left, Edit efficiencies for HEK2937^R612*/ R612*^ cells transfected with pAIO-PE2, pAIO-PE4max or PE3 plasmids. Right. Sanger traces showing a wildtype (WT), HEK293T ^R612*/ R612*^ and pAIO-PE2 edited cells. Arrow indicates the C>T SCN1A R612 location.

Next, we generated wildtype and integrase-deficient LV particles (IDLV) that could deliver the complete PE machinery. In contrast to control LV and IDLV particles that carry only the GFP gene, pAIO LV and IDLV show little GFP and marginal editing rates (1.5-1.7%) in HEK293T highlighting that further optimization for transient delivery of large LV cargo is necessary [**Supplementary data S4**].

### Removal of heterozygous *KCNQ2* R201C variant in patient-derived hiPSC

To validate the efficiency and accuracy of our pAIO-PE2-GFP and pAIO-PE4max in a more advanced model system, we tested if these systems could correct a pathogenic variant in a patient-derived hiPSC line, and compared them to the standard PE2 or PE3 systems. Nucleofected iPSCs were harvested at 72h and based on Sanger sequencing, we could not detect any edits nor indels for the CMV-PE2 and PE3 systems, which is in line with a previous study using PE3 in iPSC ^[4]^. On the other hand, pAIO-PE2-GFP nucleofected cells that had clear GFP expression after 48h **[Figure 4A]**, and pAIO-PE4max showed an average of 8% editing without additional cell sorting, and no undesired SNVs or indels **[Figure 4B**]. Because the timepoint of harvest after transfection of iPSCs differs in the literature (ranging from 48h to 96h), we then performed an experiment using only pAIO-PE4max to validate edit efficiency at 48h vs 72h vs 96h. No significant differences in edit efficiency were observed between the different time points **[Supplementary data S7]**. To find out if introducing an additional silent PAM-removing mutation could further increase editing, additional pegRNAs were designed. In contrast to our expectation, disturbing the PAM did not increase efficiency of PE at this locus [**Figure 4B**]. Interestingly, while the silent PAM removing mutation affected edit efficiency, it was virtually absent on Sanger [**Supplementary data S7]**.

**Figure 4.**
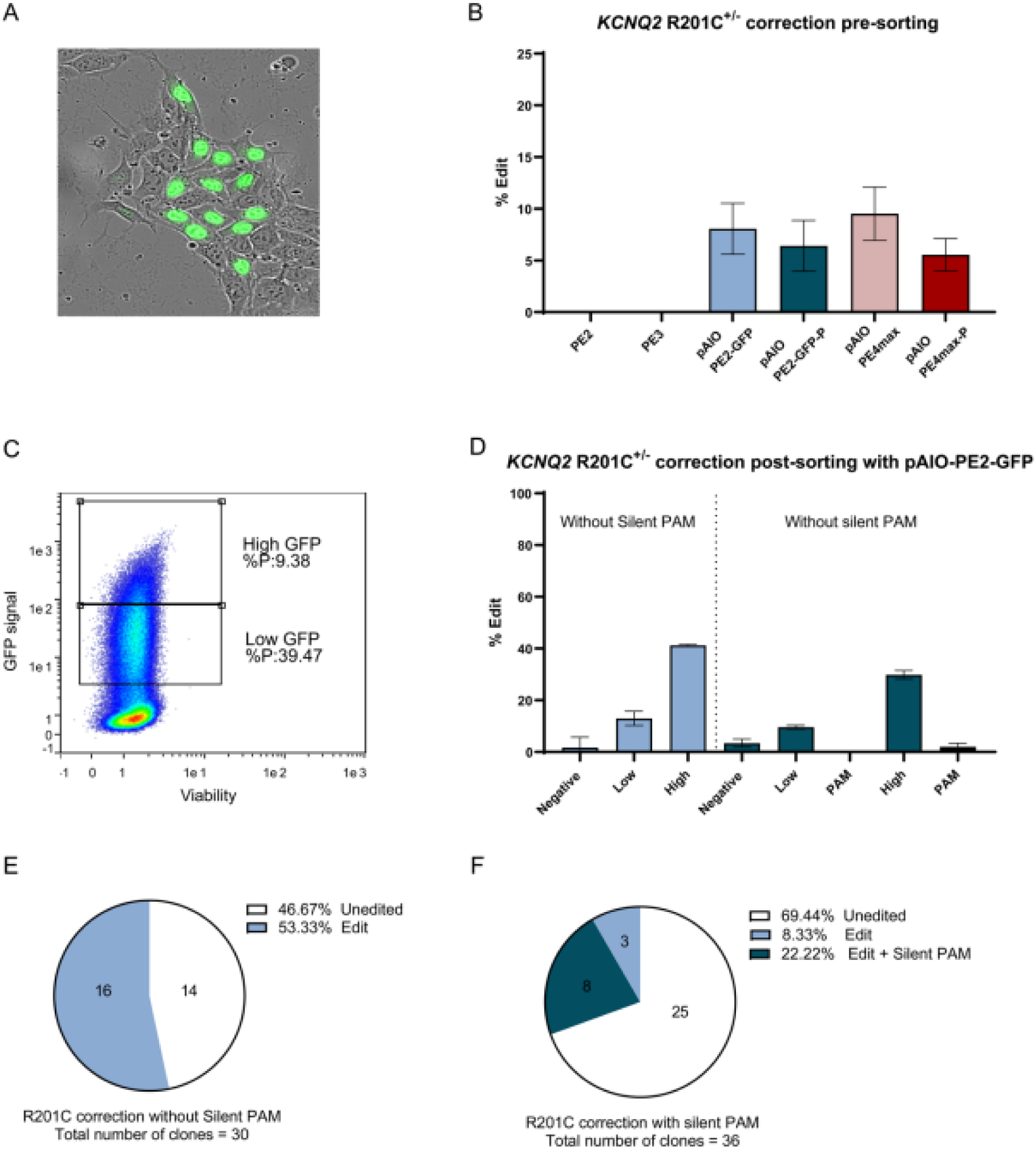
Patient-derived iPSC editing using the pAIO plasmids. A) Delivery and morphology of nucleofected iPSC. B) Correction of the heterozygous R201C mutation via pAIO plasmids compared to PE2 and PE3 with or without an additional silent PAM removing mutation. CD) Cell sorting analysis and subsequent sequencing shows increased editing if the high-GFP portion is selected. E,F) Genotyping of single-cell derived clones after pAIO-PE2-GFP nucleofection of hiPSC cells without (E) and with additional silent PAM removing mutation (F).

We next aimed to improve edit efficiency through addition of a FACS sorting step. Since pAIO-PE2-GFP and pAIO-PE4max are both effective in hiPSC with minor differences in edit efficiency, pAIO-PE2-GFP was chosen for FACS sorting due to the benefit of a GFP tag and the absence of the MLH1dn MMR inhibitor which has a risk of introducing unwanted DNA mismatches. Furthermore, we sought to investigate if the GFP signal intensity is a marker for edit efficiency. hiPSCs were sorted 48h after nucleofection. During sorting we gated the GFP population in two to separate the top half of the GFP signal (high GFP) from the bottom half of the GFP signal (low GFP signal) [**Figure 4C**]. Edit levels showed a remarkably high pAIO-PE2-GFP efficiency in the high GFP population up to 41% and a smaller increase for the low GFP population up to 13% [**Figure 4D]**. These increased efficiencies were replicated when using pAIO-PE2-GFP designed to introduce a silent PAM mutation, reaching 29% and 10% for the high and low GFP population respectively. These data show that FACS sorting on the highest GFP signal can significantly increase the edit efficiency by 5 fold, suggesting that more PE2 exposure increases editing [**Figure 4D**]. Next, using the high GFP populations, we generated clonal lines to validate edit efficiencies. For the pAIO-PE2-GFP without PAM mutation, 16 out of 30 clones (51%) were corrected [**Figure 4E]**, whereas for the pAIO-PE2-GFP with PAM mutation, 11 out of 36 clones (31%) were fully corrected [**Figure 4F**]. Interestingly, although the PAM mutations were not visible on initial sanger traces before sorting, 8 out of 11 (73%) corrected clones carried the PAM mutation. Additionally, because of the lower edit efficiency when using an additional PAM mutation, we anticipated the presence of clones with a PAM mutation but no edit, as these cells cannot be edited anymore due to the disruption of the PAM. Against our expectations no clones were found that carried only the PAM mutation, indicating a Sanger bias, delayed MMR or another unknown reason are causing this observation. Taken together, we show here for the first time that a PE2 plasmid-based system can efficiently introduce mutations in hiPSC without any selection step. We also show that pAIO-PE4max is an equally effective system. Using GFP-based FACS sorting, we could further increased the edit percentage of the pAIO-PE2-GFP nucleofected population by 5-fold. Finally, the introduction of an additional silent-PAM removing mutation was not beneficial for this locus.

### Quality control of hiPSC lines and Whole genome sequencing of corrected R201C clones

We validated 91 clones for one of the most frequently recurring CNV during iPSC culturing; the chr20q11.21 duplication. This duplication decreases apoptosis which could be favourable during subcloning iPSC lines. Using Multiplex Amplicon Quantification (MAQ) we verified that none of our clones carried the duplication **(Supplementary Data S5)**. Next, we validated in a subset of corrected clones the expression of pluripotency markers Oct4, Nanog and Sox2 **(Supplementary Data S5)**. Two corrected clones of this subset were selected to perform WGS to verify the absence of PE mediated off-target effects. WGS analysis of the two corrected hiPSC clones and the parental iPSC line shows that no alterations were made within 250 bp of all loci with homology to the pegRNA with a mismatch threshold of 4 bp. We did identify 146 and 139 novel variants respectively in the whole genome of the two corrected hiPSC clones (data not shown). These variants followed an expected distribution of intergenic:intronic:exonic variants (47:51:2), were not located in a region homologue to the pegRNA, and the majority of variants found (58%) were C:G>A:T substitutions, indicating they are probably the result of accumulation of mutations during cell culture conditions through oxidative stress mechanisms.

## DISCUSSION

In this study, we developed and tested new all-in-one (AIO) plasmids containing the full PE2 and PE4max system and showed efficient introduction or correction of Developmental Epileptic Encephalopathies (DEE) causing variants in *SCN1A* and *KCNQ2* in both HEK293T and hiPSC without off target edits.

Our experiments in HEK293T cells demonstrated efficient introduction of six different mutations in three different genes (*SCN1A, KCNQ2* and *EMX1*) with similar edit efficiencies (∼20%) using PE3. However, we observed a high yield of unintended alterations in the editing region for PE3 and other ncRNA-dependent PE systems in line with previous reports. This downside was not observed using the second-generation PE systems, PE2max and PE4max, that showed equal editing efficiency compared to PE3 cells using 3 pegRNA for 2 different loci. Due to the PE4max ability to inhibite MMR, it was previously shown to yield higher edit efficiencies compared to PE2max. Interestingly, we did not observe this for our edited loci in HEK293T cells. Because C to G conversions can be more efficiently introduced as they evade MMR, we also validated if changing a NGG motif in *SCN1A* at position c.1863-1865 (NM_001165963.4) to NCG (p.L623V) would be more efficient compared to NTG (p.L623M). No increased efficiency was observed for L623V compared to L623M using PE2max. Both observations can be explained by the MMR deficiency of HEK293T cell, which limits the MMR-inhibition effect of PE4max and the advantage of specific base changes

The few studies that tested multi-plasmid PE3 for editing in hiPSC showed a nearly undetectable edit efficiency ^[4]^, making it not suitable for the generation of iPSC-derived disease models. However, the recently published PEA1 (pAIO-PE3) all-in-one plasmid was able to increase the edit efficiency significantly in immortalized cell lines as well as mouse embryonic stem cells (mESC) when combined with a puromycin selection step ^[6]^. It did however induce similarly high unintended alterations as seen in the multi-plasmid approach. We show here the efficiency and accuracy of ncRNA-independent AIO PE plasmids, pAIO-PE2-GFP and pAIO-PE4max. To increase the functional diversity of the plasmid, we changed the CMV promotor to the elongation factor 1 alfa (Ef-1a) promoter. First, the EF1a promoter was shown to be a much stronger promotor compared to the CMV promoter in stem cells ^[7][8]^ and neurons ^[14]^. Secondly, it is one of the most stable promoters in iPSC and during *in vitro* iPSC differentiation ^[15]^. We show that our novel AIO PE plasmids, which are larger than those required for multi-plasmid PE systems, show effective delivery in both HEK293T and hiPSC cells as shown by GFP expression in 70% and 65% of cells, respectively. As expected, our Ef-1α driven AIO plasmid showed a 4-fold higher GFP expression compared to the CMV-PE-GFP in iPSC. This highlights the importance of choosing the right promoter when using plasmid-based systems and may have been a contributing factor the low success rate of previous studies using CMV-PE3 system in iPSCs ^[4]^.

We show that our pAIO-PE systems are effective for correction of the *KCNQ2* R201C mutation in a patient-derived iPSC line. Remarkably, both pAIO-PE2-GFP and pAIO-PE4max showed a correction of 8%. Although, it was expected to see an increased edit percentage in pAIO-PE4max as hiPSC have robust MMR activity ^[16]^, we saw an equal edit efficiency, which contrasts with the original report that showed a 2.5-fold increase ^[3]^. Interestingly, we observed an increased amount of cell death in the AIO-PE4max nucleofected cells, which could indicate activation of apoptotic pathways due to increased DNA damage. Furthermore, it has been shown that reduction of the MMR system in ESC due to inhibition of SIRT1 resulted in increased cell death ^[17]^. Prolonged MMR inhibition due to our plasmid approach might activate these apoptotic pathways, which might not happen when using MLH1dn mRNA as described in the original paper. Further research should be performed to clarify these discrepancies.

Our plasmid based PE2 system produced detectable editing in iPSCs without a selection procedure. To further increase edit efficiency, we sorted the pAIO-PE2-GFP nucleofected iPSC population using FACS and showed that selection of the highest GFP expressing cell population has higher edit efficiency as compared to the lower GFP expressing cell population. This shows that more copies of PE plasmid per cell, and/or a more active cell specific promoter, improves edit efficiency, probably due to the increased rate of proper transcription of the PE machinery. We found that the high GFP population gave rise to a 5-fold increase in edit efficiency (up to 41%), compared to the unsorted cells and subcloning of the iPSC line showed correction of the R201C mutation in 53% of the isolated clones. This efficient generation of subcloned iPSC lines consequently decreased the workload for colony picking and validation significantly. Finally, using WGS we could not find any genome-wide off-target effects at loci with homology to the pegRNA in two edited clonal lines, confirming prior evidence of the safety of the PE2 technology. As an outline for the future, we believe that our system might be further optimized with smaller Cas variants and cloning on a smaller plasmid backbone. We used the pLV backbone, which carries genes needed for lentiviral production, but these can be taken out if the aim is to use pDNA, downsizing the system with at least 5kb. Since >15kb pDNA with the PE machinery can be easily delivered to hiPSC with nucleofection efficiency over 60%, smaller pDNA might further improve both delivery and thereby edit efficiency. Finally, improved lentiviral production for larger cargos, ideally combined with integrase-deficient packaging plasmids to yield transient expression would enable an efficient delivery and advance translation of efficient prime editing *in vivo*.

## ACKNOWLEDGEMENT

The authors would like to thank K.Wellink, M. Olde-Nordkamp and P. van Zon for their help and advice during the project. Graphical abstract was created with BioRender.com.

---

**Supplementary S4.**
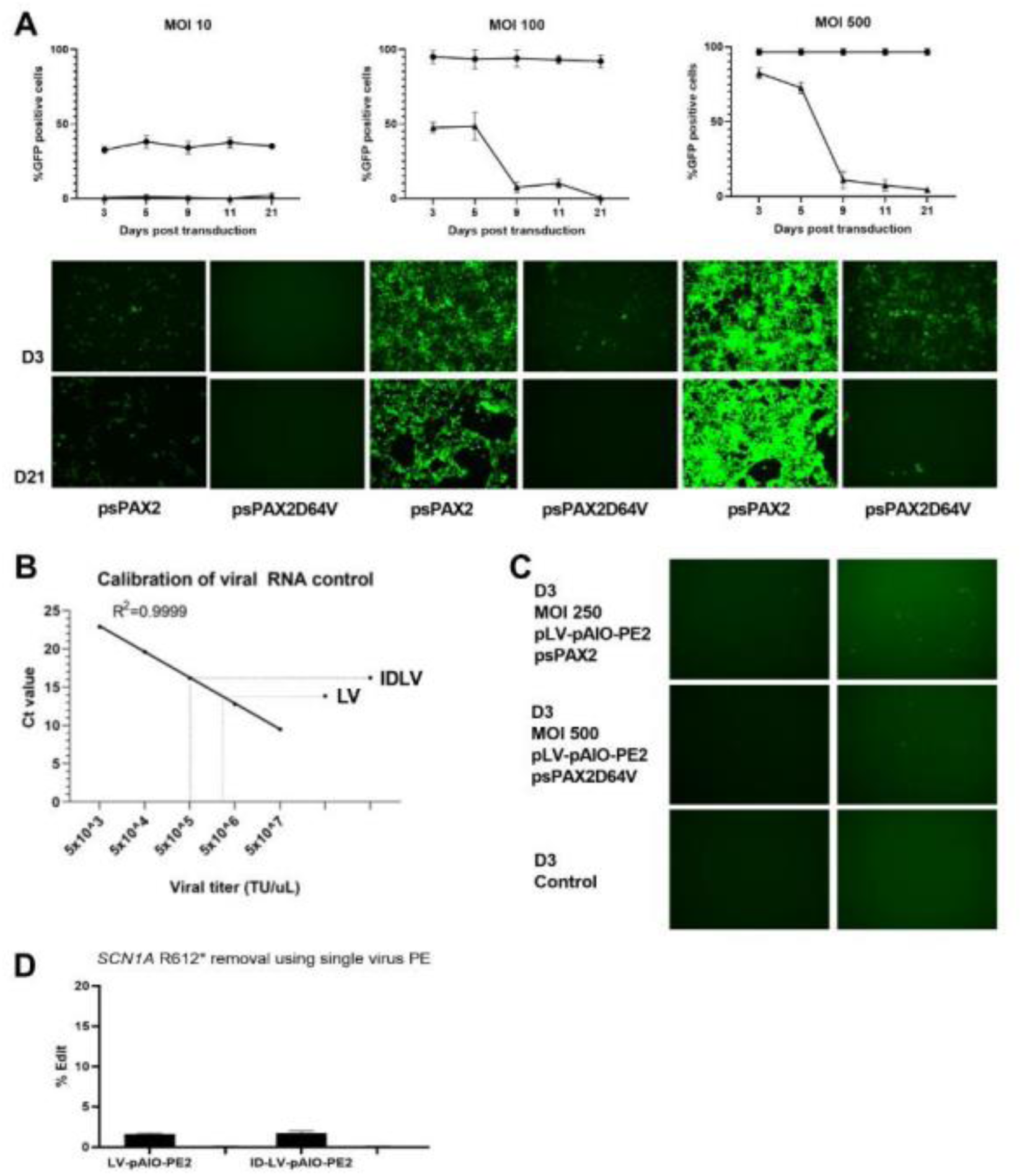
A) GFP lentivirus produced with psPAX and psPAX-D64V packaging plasmids show stable and transient GFP expression respectively, over time with various MOI. Circles; psPAX/LV, squares; psPAX-D64V/IDLV. At D21 GFP expression is absent at 10-100 MOI of IDLV indicating transient expression in contrast to stable expression of LV over time. B) Titering of pAIO-PE2-GFP LV and IDLV plotted on the control LV RNA calibration curve (Lenti-X-titering, Takara). C) Imaging of pAIO-PE2-GFP LV and IDLV at MOI 250 and 500, respectively. D) Editing rates of pAIO-PE2-GFP LV and IDLV using 250 and 500 MOI in HEK293T cells.

